# A high-throughput test of anxiety for zebrafish (*Danio rerio*) larvae: the larval diving response (LDR)

**DOI:** 10.1101/2022.05.16.492196

**Authors:** Barbara D. Fontana, Matthew O. Parker

## Abstract

**Background:** Zebrafish are used in anxiety research as the species’ naturalistic diving response to a new environment is a reliable and validated marker for anxiety-like behavior. One of the benefits of using zebrafish is the potential for high throughput drug screens in fish at the larval stage. However, at present, tests of anxiety in larvae and adults often measure different endpoints.

**New Method:** Here, for the first time, we have adapted the novel tank diving response test for examining diving behavior in zebrafish larvae to assess anxiety-like behaviors at very early-stages (7 days-post-fertilization [dpf]).

**Comparison with Existing Methods:** Current methods to examine anxiety in larvae can show low reliability, and measure different endpoints as in adults, thus calling into question their translational relevance.

**Results:** We found that 7dpf zebrafish spent more time at the bottom of a small novel tank. We validated this as anxiety-like behaviors with diazepam reducing, and caffeine increasing the time spent in the bottom of the novel environment.

**Conclusions:** This new automated and high-throughput screening tool has the potential use for screening of anxiogenic and anxiolytic compounds, and for studies aiming to understand the mechanisms underlying affective disorders.

## 1. Introduction

Anxiety exists either as a state (fear or nervousness, typically associated with a negative effect) or as a trait, in which anxiety proneness is stable across the lifespan (Hovenkamp-Hermelink et al., 2021). The emergence of anxiety traits can occur from as young as 6 years old, and can lead to an increased risk of developing subsequent substance misuse, disruptive behaviors and even suicidal behaviors (Asselmann et al., 2018). Despite pathological anxiety being common, the available drugs for treating anxiety are only palliative (i.e. not curative), and are associated with several unwanted side effects (Bandelow et al., 2017). It is therefore crucial to develop treatments high in efficacy and without significant side effects, and animal models are an essential vehicle for progressing this.

Zebrafish naturally display anxiety upon introduction to a novel environment, and novel tank tests (NTT) have been commonly used to evaluate these behaviors by exploiting the fish’s natural diving response, showing good face validity (Blaser and Rosemberg, 2012; Levin et al., 2007). These tasks also have construct validity with both anxiolytic and anxiogenic drugs reducing and exacerbating the responses, respectively (Egan et al., 2009). However, although the NTT has high reproducibility and easy validation in different laboratorial conditions (low-cost and maintenance task), its use has thus far been limited to adult zebrafish, and little is known about the diving response of younger animals.

Here, we report a high-throughput test designed for zebrafish larvae (7 days post-fertilization (dpf)), the larval diving response (LDR). The test was validated by using anxiolytic and anxiogenic drugs (diazepam and caffeine, respectively), both of which are widely used to evaluate anxiety levels in adults and zebrafish larvae (Bencan et al., 2009; Rosa et al., 2018; Tran et al., 2017).

## 2. Material and Methods

### 2.1. Animal husbandry and experimental design

Seven days post-fertilization (dpf) larval AB wild-type zebrafish (*n* = 300) were bred in-house. Eggs were collected from four tanks, and groups of ∼50 embryos were randomly assigned to petri dishes (100 × 15 mm) and reared up to 7dpf. Larvae were fed daily from day 5 (ZM000, ZM Fish Food & Equipment, Winchester, UK). Fish were kept on a 14/10-hour light/dark cycle (lights on at 9:00 a.m.; pH 8.4; 28 °C (±1 °C)). Behavioral experiments were performed between 10:00 and 16:00 h. We performed a power analysis (G* Power) based on pilot data and found an effect size of η_p_^2^ = .95 (*f =* 4.4). With power = .8 (*p* = 0.05) we required 18 zebrafish larvae per condition. As our aim here was to develop a protocol, to ensure data reliability, we ran two independent batches for habituation tests (*n =* 30 per batch) and four independent batches (*n* = 60 per batch) for the pharmacological investigation. The protocols described here are part of a preregistered study. All experiments were carried out following approval from the University of Portsmouth Animal Welfare and Ethical Review Board, and under license from the UK Home Office (Animals (Scientific Procedures) Act, 1986) [PPL: PP8708123].

### 2.2. Materials

At 7dpf, the LDR was tested using Zantiks LT automated tracking systems (Zantiks Ltd., Cambridge, UK). Caffeine (100 mg/L; Sigma Aldrich, UK) and Diazepam (5 mg/L; Sigma Aldrich, UK) were used as anxiolytic and anxiogenic drugs, respectively (Bencan et al., 2009; Egan et al., 2009; Rosa et al., 2018). In adult zebrafish, the NTT is used to measure exploration, locomotion, boldness and anxiety-like behavior (Egan et al., 2009; Levin et al., 2007). For this experiment, zebrafish larvae (7 dpf) were individually pipetted into standard cuvettes (45 mm height x 12.5 mm depth x 12.5 mm width) containing 3.5 mL of aquarium water (**Fig. 1A**). The behavioral activity was analyzed using the Zantiks LT system (Zantiks Ltd., Cambridge, UK).

**Figure 1.**
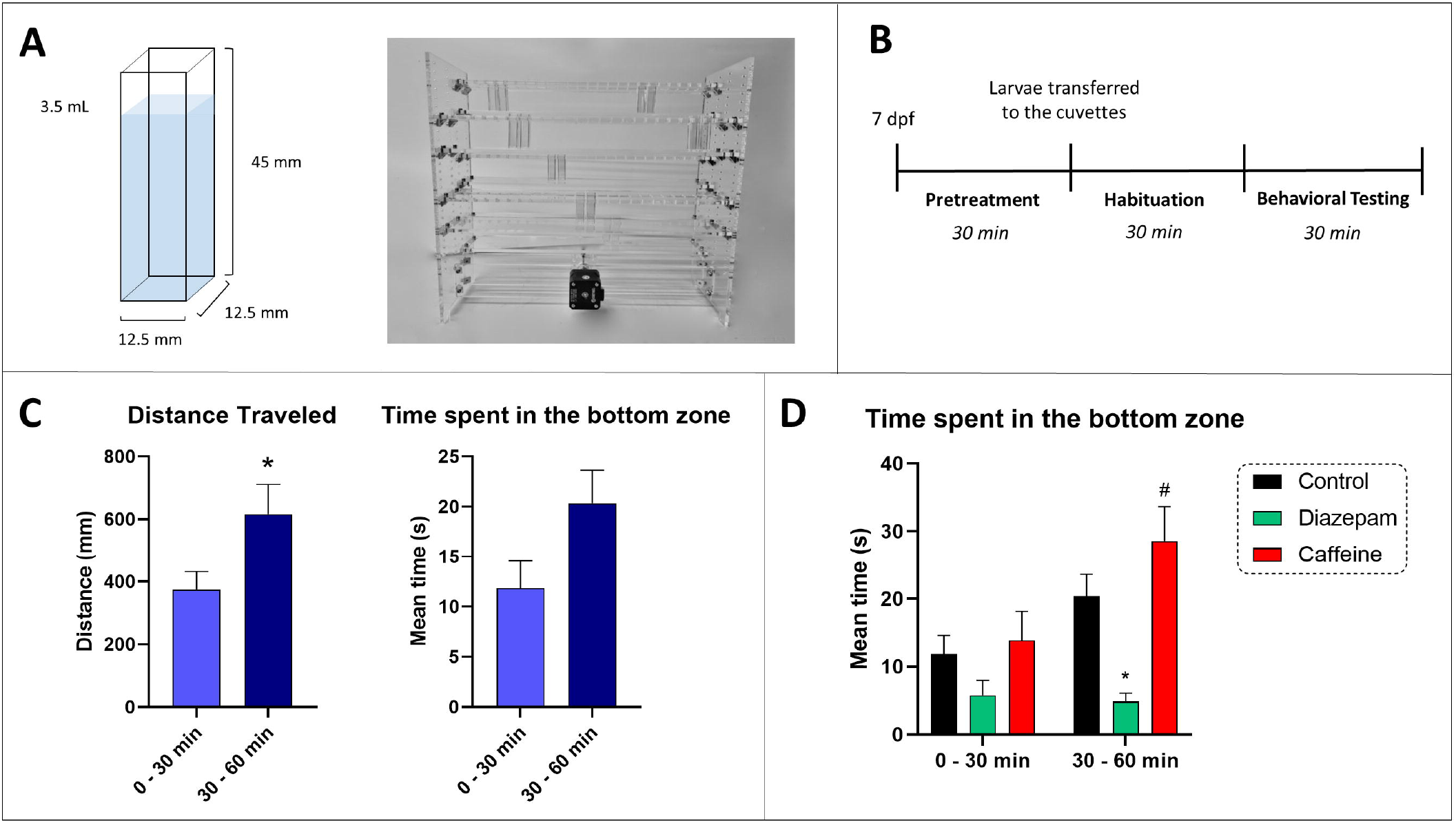
**(A)** Illustration showing the cuvette dimensions used as the behavioral apparatus for the larvae tank diving test (LDR) and a picture of the tank set up for larvae behavioral testing. **(B)** Summary of the experimental design used in this work. **(C)** Effects of time (0 – 30 *vs* 30 – 60 min) in the locomotor and diving response of larvae. **(D)** Initial data on the effects of diazepam and caffeine larvae diving response. Data were represented as mean ± S.E.M. Asterisks represent statistical differences when compared to controls of each zone (*p\** < 0.05) and octothorp represent statistical difference compared to 30 – 60 min control (*p*# < 0.05).

### 2.3. Larval Diving Response (LDR)

Zebrafish behavior in the tank diving response is typically analyzed for 5-6 min (Egan et al., 2009). Here larvae were incubated in drug solution (or fresh aquarium water) for 30 min in groups of *n* = 5 per petri-dish. They were then added to cuvettes for habituation in fresh water (see below) for 30-min, after which they were filmed for their responses for a further 30 minutes (**Fig. 1B**). These timings were determined following pilot experiments, where untreated larvae (7dpf) showed a significant increase in movement patterns only after 30 min of habituation (**Fig. 1C**). The tank was separated into three virtual areas (bottom, middle and top) to provide a detailed evaluation of vertical activity and total distance traveled (mm), number of entries, and time spent (s) in each of the tank zones. Drug response across time was also analyzed by looking at time spent in each zone across the full duration of the test (30 min).

### 2.4. Statistics

Normality of data and homogeneity of variance were analyzed by Kolmogorov–Smirnov and Bartlett’s tests, respectively. T-test was used to compare animals’ habituation to test (0 – 30 *vs*. 30 – 60 min). Two-way (3 × 2) analysis of variance (ANOVA) was used to investigate the effect of diazepam and caffeine (control. *vs*. caffeine *vs*. diazepam) in the first *vs*. last 30 min of the test. One-way analysis of variance (ANOVA) with treatment as an independent variable (three levels – control *vs*. caffeine *vs*. diazepam) was used to analyze distance travelled. Two-way (3 × 3) ANOVA was used to analyze the average number of entries and average time spent in each zone as a function of treatment (three levels – control *vs*. caffeine *vs*. diazepam) and zone (three levels – bottom *vs*. middle *vs*. top) as independent variables. Tukey’s test was used as post-hoc analysis, data was represented as mean and error of the mean (SEM), and results were considered significant when p ≤ 0.05. Because both drugs did not affect zebrafish larvae locomotion, we used ANOVA rather than ANCOVA (with distance traveled as covariate) as initially planned in the preregistration of this study.

## 3. Results

### 3.1. Larval locomotion is higher after 30 min of habituation to the environment

In **Fig. 1C** a significant increase in the distance travelled was found for control animals when comparing the initial 0 – 30 min time-period *vs*. 30 – 60 min (F _(15, 15)_ = 2.567, *p* =* 0.0309). Although time spent in the bottom was not significant for controls when comparing time periods (**Fig. 1C**; F _(15, 15)_ = 1.448, *p =* 0.0555), when looking at the habituation data comparing time-period and treatments (**Fig. 1D**), a significant effect for time-period (F _(1, 108)_ = 6.998, *p** =* 0.0094) and treatment (F _(2, 108)_ = 11.76, *p <* 0.0001) was found. Tukey′s post-hoc analysis yielded a significant difference for diazepam exposed larvae when compared to controls at 30 – 60 min (*p\** = 0.0257). Caffeine effect was also higher at 30 – 60 min, where a significant increase between caffeine 0 −30 min *vs*. caffeine 30 – 60 min was observed (*p\** = 0.0276).

### 3.2. Zebrafish larvae (7dpf) show anxiety responses which are validated by exposure to anxiolytic and anxiogenic drugs

Prior to analysis, data from 4 animals were removed due to tracking errors. In **Fig.2** the behavioral patterns of zebrafish larvae were investigated when exposed to anxiolytic (diazepam) and anxiogenic (caffeine) drugs. There were no significant changes when comparing different treatments for larvae distance travelled (F_(2,237)_ = 0.1992, *p* = 0.8195)(**Fig. 2A**). For time spent in each zone, a significant interaction between treatment and zone (F _(4,711)_ = 27.85, *p\*\*\*\** < 0.0001) and a significant zone effect (F _(3,711)_ = 74.83, *p\*\*\*\** < 0.0001) were found. No significant main effect was observed for treatment (F _(2,711)_ = 0.1228, *p =* 0.8845). There was significant difference between time spent in bottom for controls when compared to diazepam (*p** =* 0.0081) and caffeine (*p* =* 0.0115). Significant differences for time spent in the top zone were only observed for controls *x* diazepam (*p**** <* 0.0001) and no significant differences were observed for control *x* caffeine for time spent in the top zone (*p =* 0.7157). No significant differences were observed for time spent in the middle when comparing caffeine (*p =* 0.3307) and diazepam (*p* = 0.300) to control group (**Fig. 2B**). There were no differences in the number of entries (mean entries per minute) according to treatment (treatment*zone; F _(4,711)_ = 0.5241, *p* = 0.7181) or zone (F _(2,711)_ = 0.7796, *p* = 0.4590). There was a significant effect for treatment as a variable (F _(2,711)_ = 3.386, *p\** = 0.0344), but *post hoc* test showed no difference among groups, suggesting the effect was marginal (and likely unreliable as a measure) (**Fig. 2B**).

**Figure 2.**
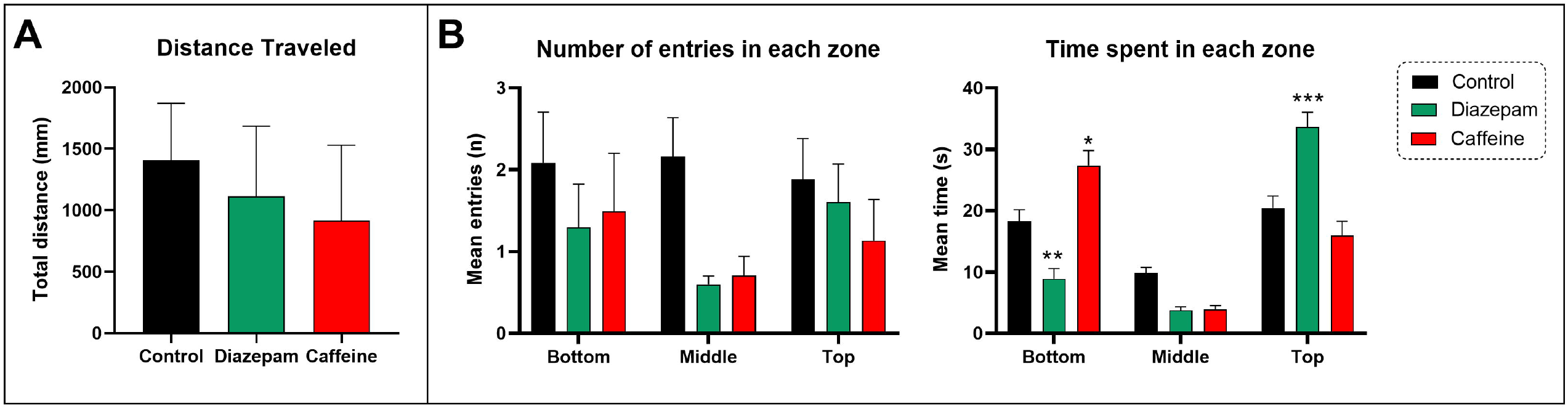
Effects of diazepam and caffeine on a larvae tank diving test. **(A)** Diazepam and caffeine did not alter larvae locomotion (*n = 80 per group)*. **(B)** Diazepam and caffeine alter diving response in zebrafish larvae by decreasing and increasing anxiety, respectively (*n = 80 per group)*. Data were represented as mean ± S.E.M. Asterisks represent statistical differences when compared to controls of each zone (*p\** < 0.05, *p\*\** < 0.005, *p\*\*\** < 0.001).

## 4. Discussion

Here, we have reported the use of a new automated method to evaluate anxiety-related behaviors in 7dpf zebrafish larvae (the Larval Diving Response: LDR) based on the NTT used in adult zebrafish. This task has good face validity, exploiting the naturalistic diving response of zebrafish. The task also has construct validity, as larvae exposed to 5mg/L diazepam (anxiolytic) show decreased anxiety-like behavior, and larvae exposed to 100mg/L caffeine (anxiogenic) show increased anxiety-like behavior while, critically, neither of the drugs affected locomotion. This is, to our knowledge, the first time that diving responses have been evaluated, quantified, and validated in zebrafish larvae, and we have demonstrated this using a fully automated, commercially available system that will allow scientists to efficiently perform high-throughput screening using larvae at an early stage of 7 dpf.

Anxiety and locomotion are often studied in zebrafish larvae. The open field test is one of the most commonly used methods, where researchers assess different parameters by placing animals in multi-well plates and recording their movements for 5 - 10 min (Ingebretson and Masino, 2013), where time spent close to the walls(thigmotaxis) is often used as an anxiety-related parameter. It could be argued, however, that thigmotaxis as a measure of anxiety in fish lacks face validity: ‘*wall hugging*’ as a protective behavior is hard to envisage in a fishes’ natural environment. In addition, despite several researchers using thigmotaxis in a multi-well plate to investigate anxiety-related behaviors in larvae, this behavioral endpoint has been criticized as being a poor predictor of anxiety in adult zebrafish (Blaser et al., 2010). For example, adult zebrafish confined in an avoidance chamber (white or black) display increased anxiety-related behaviors, but thigmotaxis and locomotor behavior were not correlated to fish avoidance (Blaser et al., 2010). Furthermore, caffeine at 50 and 100 mg/L significantly decreased time spent in the top of the NTT, indicating that those doses are anxiogenic, but thigmotaxis at both doses was not affected (Rosa et al., 2018). These data further demonstrate that thigmotaxis may not be a reliable parameter when looking at anxiety-like behaviors and that the use of complementary analysis, such as diving response may help researchers to better understand the effects of anxiogenic and anxiolytic drugs.

Zebrafish larvae develop excellent vision at 5 dpf, displaying a robust set of behaviors already at the end of the first week of development (5 – 7 dpf) (Facchin et al., 2008). Zebrafish larvae were previously shown to increase bottom dwelling (diving behavior) as an escape response to light stimulus, meanwhile, olfactory stimuli do not alter diving responses (Bishop et al., 2016). These data therefore suggest that the increased diving response to a threatful light-evoked stimulus may be an innate defense of zebrafish larvae against predation and increases in bottom dwelling could indicate an anxiogenic response. Indeed, here zebrafish showed an increased bottom dwelling when exposed to caffeine at anxiogenic concentrations, meanwhile larvae time spent in the top zone was decreased by an anxiolytic drug (diazepam) suggesting that larval diving response can be used as a parameter to measure anxiety-related phenotypes are early stages.

In the classical (adult) NTT, zebrafish spend most of their time at the bottom of the tank and gradually explore the top zone of the tank (Blaser and Rosemberg, 2012; Egan et al., 2009). Although in adults this habituation time eliminates the novel factor as a stressor and could be a limiting factor of our work, we have found the diving responses in larvae appear later (after 30-min habituation) and remain stable for ∼30 min where a higher response to anxiogenic (caffeine at 100 mg/L) and anxiolytic (diazepam at 5 mg/L) drugs were observed after this 30 min. Importantly, the use of zebrafish earlier in their lifespan has several advantages, both practically and ethically. Practically, their small size means the larvae can be screened in large numbers increasing the throughput, and the short rearing period means that multiple generations can be screened in a short period of time. As well as throughput, this has significant cost implications, with animals only being grown to a very young age therefore not requiring food and tank space. Ethically, only growing fish to a young (juvenile) age means that they are not kept in laboratory conditions beyond a few days old, thus reducing the cumulative life experience of the animals(Honess and Wolfensohn, 2010).

## 5. Conclusion

Here, we modified the NTT for adult zebrafish and designed a test to evaluate anxiety-like behavior in zebrafish larvae considering the species naturalistic diving response when facing potential threats. The method was validated using anxiogenic and anxiolytic drugs, where 5 mg/L diazepam reduced, and 100 mg/L caffeine increased anxiety-like behaviors in 7dpf zebrafish. This new fully automated, commercially available, high-throughput screening method could accelerate the discovery of new drugs for treating neuropsychiatric disorders that affect anxiety.

## Acknowledgments

This study was supported in part by the CAPES (Brazil) - Finance Code 001 at the University of Portsmouth, UK (BDF). MOP also currently receives funding from the Alzheimer’s Research UK, INTERREG (EU), Dstl (UK), and NC3Rs (UK). The funders had no role in study design, data collection, and analysis, decision to publish, or preparation of the manuscript.

## Conflict of Interest

The authors declare no conflict of interest.

## Open Practices Statement and Data Availability

The data for all experiments are available at (https://osf.io/kpmvh) and more info about the study preregistration can be found at https://osf.io/qxpu8.

## Notes

### Competing Interest Statement

The authors have declared no competing interest.

